# CODI: Enhancing machine learning-based molecular profiling through contextual out-of-distribution integration

**DOI:** 10.1101/2024.06.15.598503

**Authors:** Tarek Eissa, Marinus Huber, Barbara Obermayer-Pietsch, Birgit Linkohr, Annette Peters, Frank Fleischmann, Mihaela Žigman

## Abstract

Molecular analytics increasingly utilize machine learning (ML) for predictive modeling based on data acquired through molecular profiling technologies. However, developing robust models that accurately capture physiological phenotypes is challenged by a multitude of factors. These include the dynamics inherent to biological systems, variability stemming from analytical procedures, and the resource-intensive nature of obtaining sufficiently representative datasets. Here, we propose and evaluate a new method: Contextual Out-of-Distribution Integration (CODI). Based on experimental observations, CODI generates synthetic data that integrate unrepresented sources of variation encountered in real-world applications into a given molecular fingerprint dataset. By augmenting a dataset with out-of-distribution variance, CODI enables an ML model to better generalize to samples beyond the initial training data. Using three independent longitudinal clinical studies and a case-control study, we demonstrate CODI’s application to several classification scenarios involving vibrational spectroscopy of human blood. We showcase our approach’s ability to enable personalized fingerprinting for multi-year longitudinal molecular monitoring and enhance the robustness of trained ML models for improved disease detection. Our comparative analyses revealed that incorporating CODI into the classification workflow consistently led to significantly improved classification accuracy while minimizing the requirement of collecting extensive experimental observations.

**SIGNIFICANCE STATEMENT:** Analyzing molecular fingerprint data is challenging due to multiple sources of biological and analytical variability. This variability hinders the capacity to collect sufficiently large and representative datasets that encompass realistic data distributions. Consequently, the development of machine learning models that generalize to unseen, independently collected samples is often compromised. Here, we introduce CODI, a versatile framework that enhances traditional classifier training methodologies. CODI is a general framework that incorporates information about possible out-of-distribution variations into a given training dataset, augmenting it with simulated samples that better capture the true distribution of the data. This allows the classification to achieve improved predictive performance on samples beyond the original distribution of the training data.

## INTRODUCTION

Technological advances in molecular analytics increasingly enable the probing of biological systems. Distinguishing between physiologically relevant states from quantitative molecular fingerprints presents a new opportunity for *in vitro* phenotyping. Extensive efforts are thus dedicated to developing standardized procedures involving streamlined biological sampling, post-collection handling, and sensitive quantitative measurements. Nevertheless, empirical datasets remain susceptible to diverse sources of variability, both analytical and inherently biological [1–9]. Obtaining a dataset that reflects a realistic data distribution is often resource-intensive, costly, and, in some cases, impossible. This applies especially in the context of clinical studies, covering all pathophysiological strata, studying rare disease, or longitudinally probing the same system over time. Exploratory studies are thus often limited in size and scope, making it challenging for a given “training” set to be representative of the true unseen “test” domain. Consequently, when applying a developed machine learning (ML) model to independently collected and experimentally measured samples, the model may fall short of achieving the expected efficacy [8, 10–14].

While traditional approaches often rely on standardizing experimental workflows and creating computer-aided processing techniques to reduce unwanted empirical noise [6, 15–19], complete noise removal is likely unattainable. Failure to account for noise and distributional shifts may mask the true biological patterns of interest, potentially misleading an ML algorithm into utilizing confounding information that is unlikely to be reproduced. This failure is due to violating the assumption underlying (supervised) ML algorithms that the training and testing data are independent and identically distributed (i.i.d.) [10, 11, 20–22]. To decode the information that a collected dataset may hold, understanding and accounting for analytical and biological variability is critical to ensuring successful model generalization.

The concept of out-of-distribution (OOD) generalization has very recently garnered attention in ML research to address the shortcomings of i.i.d. assumptions [20, 23, 24]. This paradigm shift acknowledges the unpredictability of unseen data, prompting exploration into methods that better accommodate distributional shifts to generalize beyond the training set. OOD generalization has been extensively explored in computer vision and natural language processing tasks [23–26]. However, there is a critical lack in the development of OOD generalization techniques in molecular analytics involving vibrational spectroscopy, NMR spectroscopy, and mass spectrometry, as well as in clinical chemistry analytics.

To address these challenges, here we develop and empirically test a hybrid experimental and computational modeling framework. We explore OOD generalization in the context of molecular analytics and propose to recognize the variations that arise from the analytical procedure as integral components of real-world observations (Fig. 1). We introduce Contextual Out-of-Distribution Integration (CODI), a modeling framework that paradoxically embraces measurement variability and inherent complexities of biological systems, transforming them into valuable properties that can be utilized. CODI first involves experimental data to evaluate their distributional characteristics. Following the characterization, we deliberately introduce these distributional characteristics into a studied, independent, dataset through the *in silico* generation of synthetic data. These synthetic data mimic the system(s) of interest, while expanding the distribution of the original seed data to incorporate information about sources of variance that were crucially OOD and missing.

**Figure 1.**
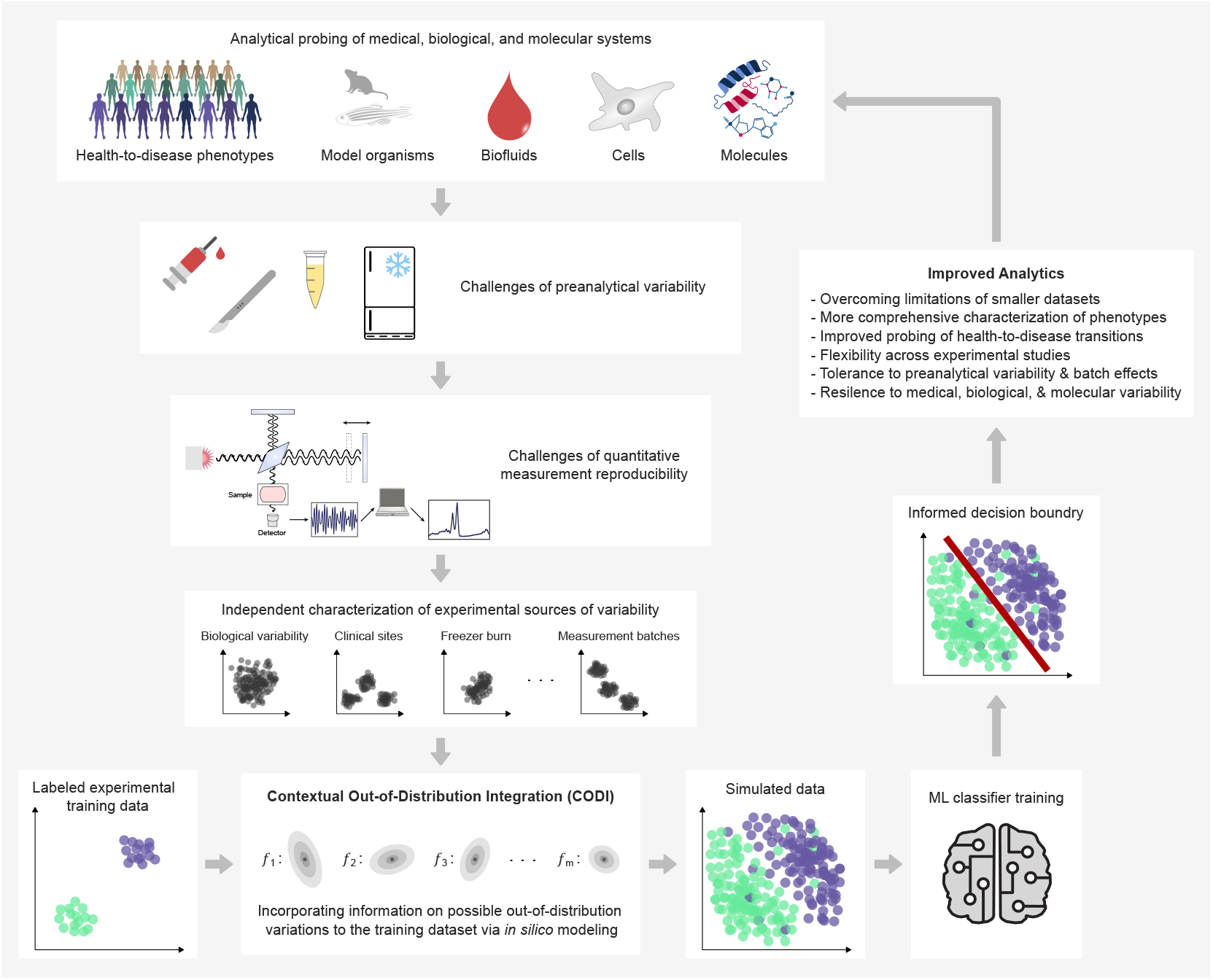
Overview of problem context and CODI’s methodology. Given a biological, medical, or molecular system of interest, we are presented with a task of classifying distinct groups of samples (e.g., healthy vs. diseased observations). However, variations stemming from several biological and (pre)analytical aspects in the empirical workflow may impact captured measurements in different ways. CODI leverages independently characterized sources of variability and incorporates them into a labeled training set of experimental observations. This process generates a larger set of simulated samples, with a more representative data distribution. Training an ML classifier on simulated samples enables it to learn a decision boundary that separates classes of samples in a more informed manner, increasing the likelihood of generalizing to unseen test samples. This approach enables sample characterizations that are more robust to variations in the empirical workflow.

To establish the concept and evaluate it in a realistic setting, we apply our method to experimental infrared (IR) spectroscopic data to aid *in vitro* blood-based diagnostics. The advantage here lies in cross-molecular fingerprinting, where quantitative analytical measurements capture the breadth of changes in the molecular landscape of complex samples as indicators of systemic health and disease. We test our method in the framework of three independent longitudinal clinical studies spanning up to an 8-year follow-up period [27–29], as well as a case-control study to detect four common cancers [30]. Our results demonstrate that integrating CODI into the classification pipeline enables the creation of more representative datasets, arbitrarily large in size, that empower ML algorithms to more effectively capture reproducible signals in biological datasets. Ultimately, we showcase how the proposed framework leads to significantly improved classification output on unseen, independently measured test samples, ensuring robust predictions despite shifts in data distribution.

## RESULTS

### Characterizing empirically-observed variability

We previously introduced an *in silico* model that generates 1D spectra of complex biological samples, focusing on IR absorption spectra [31]. Our initial work explored the impact of varying levels of between-person biological variability on classification efficiency in simulated case-control conditions. Building on this foundation, we extend the model beyond the theoretical framework. We generate data simulating longitudinal and/or case-control settings, accounting for diverse sources of possible variability that we experimentally characterize. To assess its practical applications, we explore the capacity of the modeling framework to computationally generate larger training sets that are more robust to biological and analytical variabilities.

CODI is a relatively simple statistical procedure that relies on the characterization of data distributional characteristics. In essence, we capture the differences between sets of experimental observations (***u***_*i*_ − ***v***_*j*_), henceforth called calibration measurements (e.g., a control sample repeatedly measured under varying laboratory conditions). By selecting calibration measurements from a given measurement pool that exhibit a level of variation, we can scale the differences by a random variable assuming a probability distribution, and combine them over the whole set of utilized calibration measurements. This aggregation may then be added onto an independent experimental measurement ***x***_*k*_ to create a new simulated measurement. Such a simulation approach can be repeatedly applied to generate a cohort of measurements in arbitrary size. The generated cohort, as a whole, would reflect the variability properties observed between ***u***_*i*_ and ***v***_*j*_ onto ***x***_*k*_. If the set of calibration measurements would reflect a new source of variability that was unobserved in a given training set of measurements {***x***_*k*_ | *k* = 1, …, *n*}, a new level of variability would be introduced onto the training set of measurements. This strategy allows for the creation of realistic synthetic data without requiring the adjustment of any free parameters controlling the data generation – other than the number of generated observations. A detailed description of the modeling approach is provided in the Methods and Supplementary Information.

In our example applications of CODI, we introduced several distinct sets of calibration measurements to model different sources of variability that may be observed in IR spectral measurements of blood-based media (Fig. 2**a**). Within these calibration measurements are characteristics of empirical variability stemming from inherent biological factors, variations in sample collection and handling, as well as instrument-specific measurement noise and drifts (Methods, Supplementary Information).

**Figure 2.**
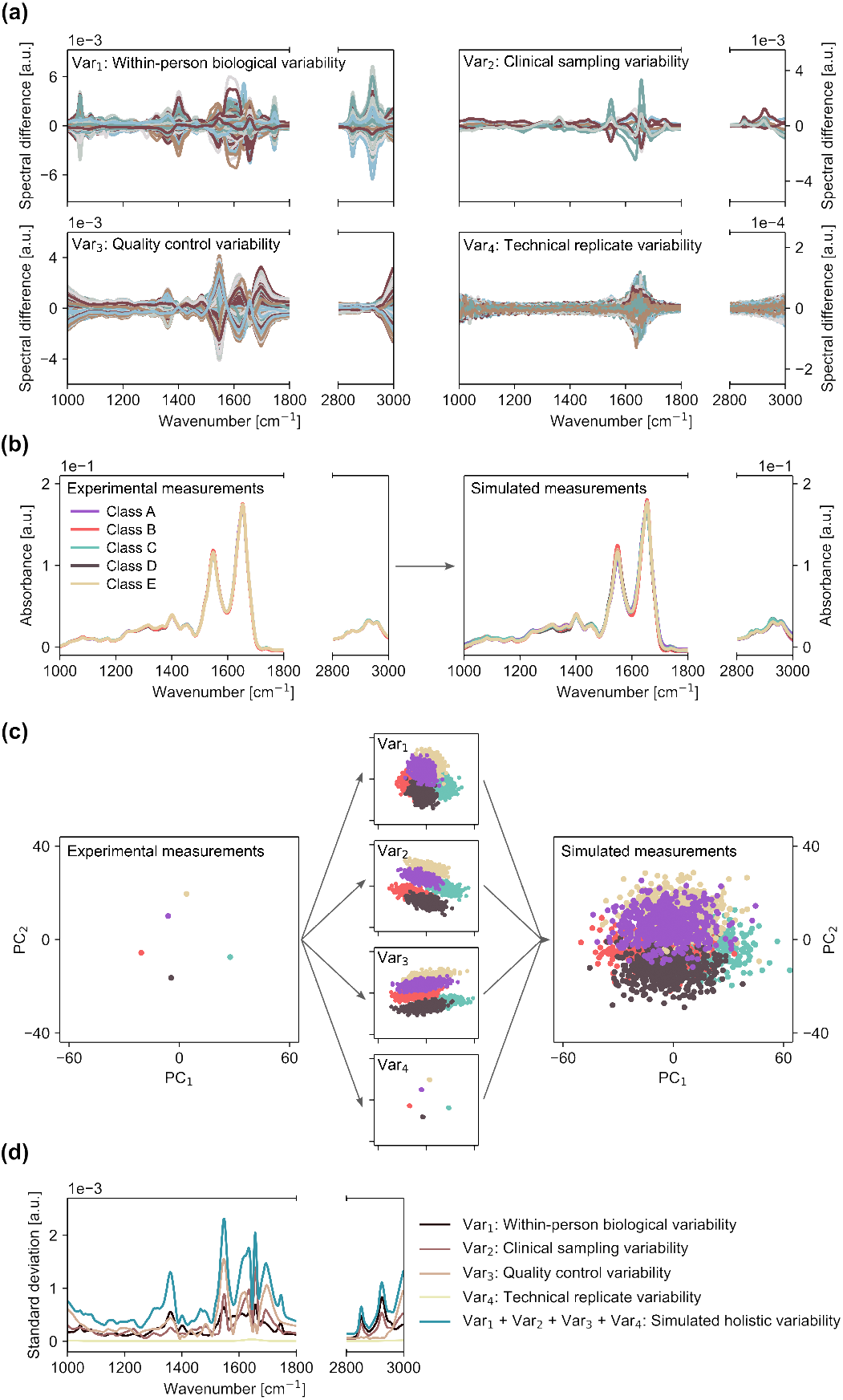
Applying CODI to introduce measurement variability onto illustrative experimental IR spectra. (**a**) Four distinct sources of possible variability were characterized from calibration sets of experimental observations. (**b**) By applying CODI, the four sources of variability were introduced to five illustrative experimental blood-based spectra (left panel) to generate a larger set of simulated spectra (right panel). (**c**) Principal component analysis (PCA) on the five illustrative experimental spectra (left panel), on a larger simulated set of spectra that introduced only one out of the four characterized sources of variability (middle panel), and on a simulated set of spectra that introduced all four sources of variability to the five illustrative experimental spectra (right panel). (**d**) Comparison of the standard deviation across the fingerprint spectral range for each characterized source of variability and the standard deviation of simulated measurements (incorporating all four sources of variability), resulting in an overall increased standard deviation.

When addressing biological variability, the calibration measurements ***u***_*i*_, ***v***_*j*_ are selected to be experimental measurements of the same individual over time, capturing a level of within-person biological variability (Fig. 2**a**, upper left), as defined previously [29]. Alternatively, opting to set the calibration measurements to be of different individuals would yield a level of between-person biological variability, as demonstrated previously [31].

Further variations that stem from different clinical sample collection sites (e.g., clinics), clinical study protocols, and sample handling procedures may be effectively represented by selecting calibration measurements characteristic of samples derived from different clinical studies (Fig. 2**a**, upper right). The same concept can be extended to model realistic variations that arise from experimental procedures like sample storage temperature and duration, aliquoting procedures, and measurement device drifts. For example, quality control (QC) samples, may be subjected to diverse handling and storage conditions. Performing measurements of QCs under different operating conditions for the measurement device, including instances of recalibration, routine maintenance, or changes in the surrounding environment would enable the QC measurement dataset to mimic potential variations in both laboratory procedures and instrumental drifts (Fig. 2**a**, lower left). Further, independent, measurements of technical replicates (e.g., pure water) performed over extended periods can facilitate a clearer distinction between instrumental device noise and laboratory variations (Fig. 2**a**, lower right).

The overarching goal of characterizing diverse sources of variability is to realistically simulate the data distribution that may be encountered in the empirical workflow. It is crucial to recognize that this is achieved through the utilization of measurements that are independent of the original training set and unrelated to the specific questions posed by it (i.e., class-invariant). Therefore, the characterized source of variability can be repeatedly used across a diversity of classification tasks. When extending the concept to other molecular systems or measurement modalities, similar calibration sets may be adapted to characterize the variations relevant to the studied conditions.

### Introducing experimental variability *in silico*

As an illustrative example, we applied the CODI modeling framework to five experimental spectra of blood plasma spectra to generate a larger set of simulated measurements that reflect an increased level of variance (Fig. 2**b**). The five original spectra may be considered to be a training set, with each measurement representing a labeled class (Fig. 2**b**, left). Using the five measurements as a seed input, the CODI framework allowed us to generate a larger and more representative training set of measurements (Fig. 2**b**, right).

Principal component analysis (PCA) applied to the original and simulated measurements reveals that each source of measurement variability affected the spatial distribution of the seed data differently across the first two components (Fig. 2**c**). In other words, each set of calibration measurements – modeling different variability properties – affected linearly independent data features. Once all four sources of measurement variability were incorporated into the original seed data (Fig. 2**c**, right), the simulated measurements occupied a larger cloud of data points, while still maintaining their distinct cluster centroids.

Similarly, an examination of the standard deviation between each source of modeled empirical variability and the simulated measurements reveals that the standard deviation of simulated measurements was higher than that of each independent source of empirical variability (Fig. 2**d**). While it may seem counterintuitive that a simulated dataset with increased variance could offer added value compared to the existing experimental observations, this variance contains valuable, usable information. The principle relies on the assumption that the simulated measurements represent unaccounted fluctuations in a smaller set of measurements, out-of-distribution, that are likely to occur when a larger set of experimental observations is available.

So far, we have shown how CODI can be used to introduce additional sources of variability into an existing measurement dataset to enrich its information content. The value of the method for real-world applications is examined in the following sections.

### Application to longitudinal study settings

In longitudinal clinical studies that involve the collection of biological specimens tainted by attrition and loss-to-follow-up over time, great efforts are required to gather sufficiently large datasets. Typically, individuals participate in an initial baseline sample collection, followed by extended waiting periods for subsequent collections from the same individuals. In situations where only few samples are initially available per individual, the challenge arises in extrapolating meaningful insights to later collected and measured follow-ups – owing to the dynamic nature of the empirical procedure as previously described. To examine whether our proposed approach offers added value when severely limited samples are available for analysis, we employ the CODI framework in the context of longitudinal analyses (Fig. 3).

**Figure 3.**
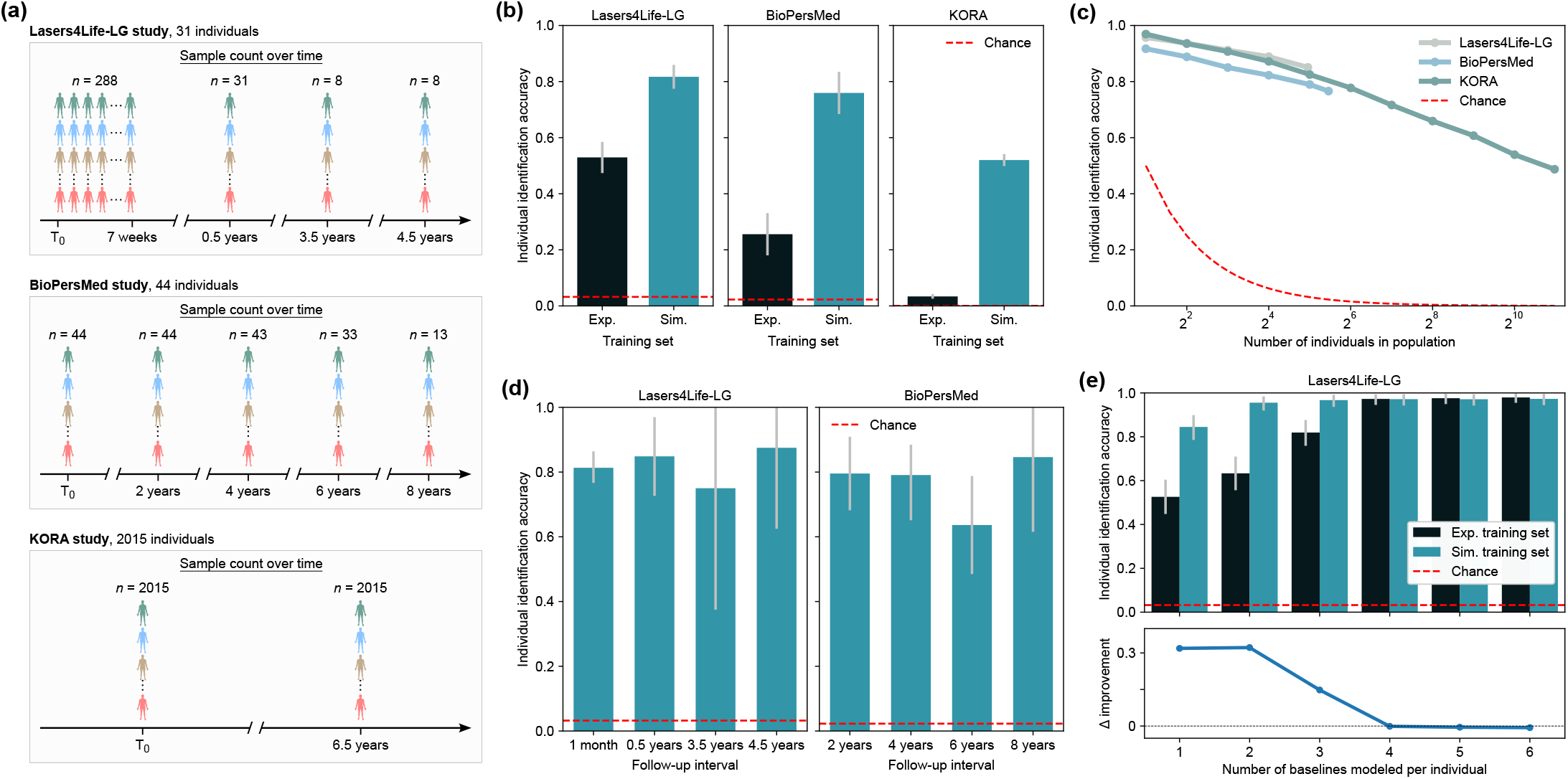
CODI enhances personalized fingerprinting through more accurate long-term molecular profiling. (**a**) Setup of three independent longitudinal clinical studies in which same individuals repeatedly participated in venous blood sampling over time. Experimental IR spectroscopic measurements were performed on blood plasma and utilized in this analysis. (**b**) Individual identification efficacy utilizing only a single baseline IR measurement per individual across the three study cohorts. Black bars depict the classification accuracy using experimental (“Exp.”) baseline measurements for training. Blue bars depict accuracy using simulated (“Sim.”) measurements generated by CODI using a baseline IR measurement per individual as a seed input. Classifier testing was performed on the same experimental follow-up measurements of each individual for both training approaches. (**c**) Dependence of identification accuracy on number of individuals in the populations (i.e., number of classes). Individuals were randomly selected, at varying population sizes, and CODI was applied using a single baseline IR measurement per individual as a seed input. Classifier testing was performed on experimental follow-up measurements of each individual included in the training set. (**d**) Dependence between the identification accuracy and the follow-up time axis on applications involving the Lasers4Life-LG and BioPersMed cohorts. (**e**) Modeling an increasing number of experimental baseline measurements per individual as a training seed. Blue bars depict the identification accuracy on simulated training sets, while black bars depict the identification accuracy on experimental training sets. Model testing was conducted on the same experimental follow-ups beyond the first 6 baseline measurements per individual. The lower panel illustrates the difference between accuracy observed from the experimental and simulated training sets.

We first considered three independent clinical studies that followed individuals over extended periods (Fig. 3**a**). The Lasers4Life-LG study cohort [29] comprised of 31 individuals that repeatedly donated blood samples at irregular follow-up intervals. The study commenced with a 7-week baseline monitoring period, during which 288 samples were collected through repeated donations. The initial baseline donation period was followed by three additional donations, spanning up to 4.5 years, during which one sampling point was considered per individual. In the BioPersMed study [27], a subcohort of 44 individuals repeatedly participated over an 8-year follow-up period, with a 2-year interval between each donation. In the KORA study [28], a subcohort of 2015 individuals participated in two donations, separated by a 6.5-year follow-up interval. Blood plasma was processed from all samples and measured via absorption IR spectroscopy (Methods).

### Empowering long-term molecular profiling

The concept of identifying individuals based on different biofluids has been demonstrated with IR spectroscopy, NMR spectroscopy, and mass spectrometry as fingerprinting modalities [29, 32–34]. We previously demonstrated that plasma- and serum-based IR fingerprints can identify individuals over a span of 6 months [29]. Such an application inherently relies on the stability of measurements over a study period, often requiring the comparison of measurements acquired at different times despite inevitable experimental drifts [35]. In previous works [29, 32–34], several measurement data points from the same individuals were required to adequately train a multi-class classifier to identify individuals (typically 8-42 samples).

We set out to test the possible value of our CODI framework for enhancing longitudinal studies, using the individual identification task as a readout metric. To test the limits of the framework, here we considered a scenario with severely limited training data – relying only upon a single experimental observation per class for training, i.e., one measurement per individual (Fig. 3**b**). We utilized the first baseline measurement of each individual to train a multi-class classifier to identify individuals from their follow-up measurements. We then compared this prediction efficiency to a classifier trained on a simulated set of measurements created through the CODI framework that used the experimental baselines as initial seed data. Within the CODI framework, we modeled the four previously described sources of variability (Fig. 2**a**), which were characterized from data sources independent of the experimental seed data (Supplementary Information). This allowed us to generate 1000 simulated measurements per individual that were used for training the classifier.

This investigation revealed that the classifier trained on simulated measurements had a remarkably improved prediction efficiency compared to the classifier trained on experimental measurements (Fig. 3**b**). Across the three cohorts, the individual identification accuracy improved from 0.53 to 0.82, from 0.26 to 0.76, and from 0.03 to 0.52. This demonstrated that the (informed) incorporation of data variance was indeed capable of enabling the classifier to better generalize to unseen test samples. To examine which sources of data variability aided the most in boosting the classification efficiency, we re-performed the above analysis, but systematically eliminated one of the four sources of variability incorporated in the CODI modeling framework (Supplementary Fig. S1). We found that the within-person biological variability over time and measurement variability of the same quality control samples over the course of measurements were the most critical contributors for the success of the classification. Including the variability stemming from technical replicates and clinical sampling had minimal impact on the classification, compared to the two aforementioned sources of variability. This highlighted the importance of incorporating information on the data distribution that was truly missing from the original experimental training set – not just the incorporation of (random) added variance.

It is crucial to recognize that the population size varied between the cohorts. The decreased accuracy observed in the KORA cohort (involving 2015 individuals) is thus not directly comparable to that of the Lasers4Life-LG cohort (involving only 31 individuals). This is due to the fact that the more individuals exist in a dataset, the more likely it is that their fingerprints will overlap with one another – making the task of identifying individuals more challenging. Despite this, it was very surprising and encouraging to observe that nearly half of 2015 individuals can be identified from IR molecular fingerprints when combined with the proposed modeling approach – and requiring only the venous blood sampling of only a single baseline sample per individual.

Observing that the identification accuracy decreased with an increasing population size prompted us to further investigate this dependency (Fig. 3**c**). We first trained a classifier on simulated measurements utilizing the first experimental baseline measurements of only 2 individuals and, as previously, tested on their follow-ups. This procedure was repeated several times, using 2 other randomly selected individuals. We then performed the same procedure, but on 4, 8, 16, and so on individuals. This analysis revealed that the identification accuracy, depending on the population size, follows a nearly perfect logarithmic trend. This intriguing finding draws from information theory and may be explored further to quantify the informational content of diverse molecular fingerprints. Remarkably, these results were reproducible on three independent cohorts, revealing that a similar identification accuracy can be achieved when the datasets involve similar population sizes.

In the above application, follow-up measurements of all individuals were pooled together and the accuracy of identification was averaged, independent of the follow-up time axis. This prompted the question of whether the individual identification accuracy was dependent on the time interval between follow-up measurements and the baseline. In other words, is it more difficult to identify an individual 8 years after their baseline sample was assessed than from a 2-year follow-up? To investigate this, we grouped the follow-up measurements by their time differences to the baseline and examined whether any temporal trend was observed in the identification accuracy (Fig. 3**d**). This analysis was only possible on the Lasers4Life-LG and BioPersMed cohorts, since the available KORA cohort only involved one follow-up. Here, we revealed that the identification accuracy did not depend on how far off the follow-up was from the baseline. Very surprisingly, the accuracy remained relatively stable even over an 8-year follow-up period. Although it is crucial to recognize that the number of test samples in the later follow-up years was limited (Fig. 3**a**), this is the very first experimental result over such long-lived fingerprint stability.

Altogether, the above investigations were made possible by using the CODI framework which enabled applications that were previously unfeasible with limited experimental observations.

### Comparison to domain-agnostic augmentation schemes

The CODI modeling framework inherently relies on *a priori* information on potential sources of measurement variability. In contrast, domain-agnostic augmentation methods employ generic transformations on input seed data to simulate new observations (e.g., introducing additive or multiplicative noise). By eliminating the need for *a priori* information, such a strategy is simpler to implement than CODI. To examine whether CODI yielded an advantage over other augmentation methods, we reperformed the above analysis by applying several methods of augmenting the spectral measurements (Supplementary Fig. S2). We found that the CODI method of generating training sets significantly outperformed all other methods of data augmentation. This underscored the value of incorporating contextual *a priori* information into the data augmentation process to enable the classification to generalize beyond the original training set in an informed manner.

### Personalized multi-baseline modeling

In the above quest of examining the value of the CODI framework, we relied on a single baseline measurement per individual. We then questioned to what extent can the classification be made more robust when more training instances per class are available. Specifically, considering that a single baseline measurement may be an outlier, we examined the dependence between the number of training instances per individual and the identification accuracy (Fig. 3**e**). For this analysis to be properly investigated, individuals would have to be repeatedly sampled in a given baseline monitoring period. Among the three clinical studies, only the Lasers4Life-LG study facilitated such a setup [29].

Here, we utilized data from up to the first 6 baseline measurements per individual from the Lasers4Life-LG cohort to be used as an experimental seed for training. Then, we simulated a training set that consisted of 1000 measurements per baseline and investigated how the identification accuracy depended on how many experimental baselines were modeled per individual. Classifier testing was performed on the remaining follow-ups that were beyond the first 6 baselines of each individual. This investigation revealed that the identification accuracy following the simulation-based approach could indeed be improved when more than one baseline measurement was modeled per individual (Fig. 3**e**, blue bars). When using only one baseline measurement, an accuracy near 0.85 was achieved. Surprisingly, including only one additional baseline already led to an improvement of a nearly perfect prediction efficiency, achieving an accuracy of 0.96.

As a comparable benchmark, we again examined the dependence between the identification accuracy and the number of baselines modeled per individual, but now training the classifier directly on the experimental measurements (Fig. 3**e**, black bars). The classifier was first trained on one experimental baseline per individual, then again on two, and up to 6 baselines of each individual. Testing was performed as previously - on the remaining follow-ups beyond the first 6 training baselines per individual. Here, it was again revealed that the simulation-based approach had a significant advantage over the experimental modeling approach - but only when few observations per class were available (≤ 3 baselines per individual). Once ≥ 4 experimental baselines per individual were available for training, the experimental modeling approach had also achieved a near perfect prediction efficiency and thus no advantage was seen by applying CODI. This underscored the impact of our proposed modeling paradigm in contexts with only limited experimental datasets. Once sufficiently large experimental datasets are available, the simulation-based training approach may not provide an advantage over training directly on experimental data.

Altogether, these findings show that the CODI framework can enable the establishment of a more reliable “baseline” per individual – one that is more resilient to analytical and biological variations and can more robustly enable ML generalization. We further demonstrate that IR molecular fingerprints are highly stable and individual-specific. Previously, this was only demonstrated on the time-frame of 6 months [29]. In the current study, we extend these findings to a medically relevant time frame of 8 years. These results form the foundation for future applications of blood-based IR fingerprinting as a modality of personalized monitoring of human health over time, potentially requiring as few as one or two baseline samplings.

### Cross-specimen generalization

Molecular profiling applications involve the use of diverse sample specimens - e.g., serum or plasma as cell-free products of systemic blood (Fig. 4**a**). Selecting an appropriate specimen is typically made in a study design phase, considering factors like ease of collection and biological relevance [36, 37]. However, limitations may arise from a preemptive selection as insights gained from one specimen may not generalize when transferred to another. For instance, assume a dataset of plasma spectra is available. Later, the need arises to classify and compare unlabeled spectra that originate from serum samples. This prompted an intriguing question: how well would a classifier trained on plasma spectra perform when tested on serum spectra? The straightforward answer is that the classification is likely to fail, due to underlying molecular differences between the specimens [38]. Effective classification necessitates the inclusion of training instances from different specimens, each with sufficient representation to capture class-specific distributions – a highly resource-intensive process. As proof-of-principle, here we demonstrate the potential versatility of the CODI framework to enable such an application, while minimizing the need for extensive biological dataset collection.

**Figure 4.**
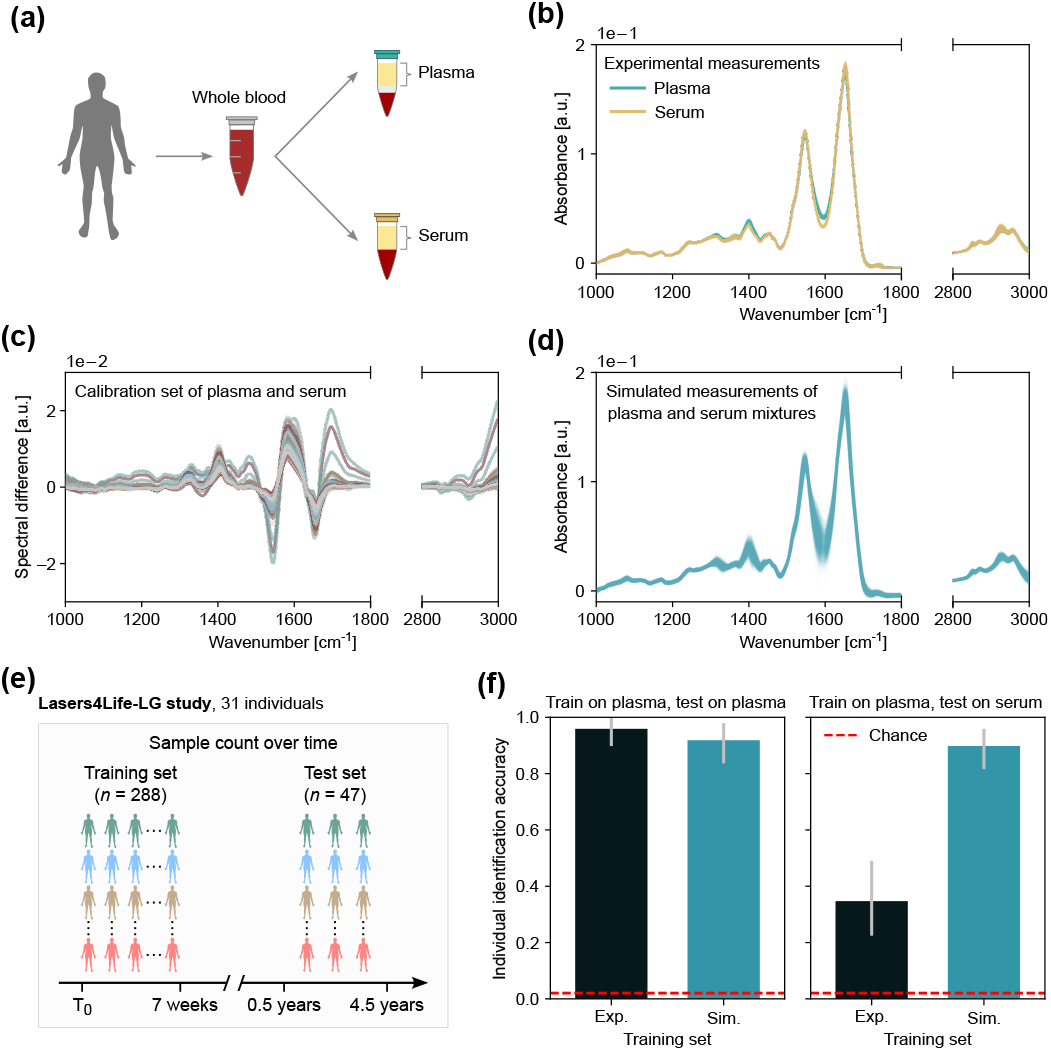
CODI enables classification flexibility across biological specimen variants. (**a**) Plasma and serum were collected at the same sampling point as cell-free products of whole venous blood. (**b**) Experimental spectra were measured from several plasma and serum samples of the same individuals. (**c**) Differences between spectra of plasma and serum, processed from the same whole blood sample, were calculated to reveal the characteristic variations between the specimens. (**d**) CODI framework enabled the creation of simulated spectra of plasma/serum mixtures by utilizing the characteristic variations between the specimens as a calibration set. (**e**) Setup of Lasers4Life-LG cohort in which the same individuals repeatedly participated in venous blood sampling over time. Donations were split into a training set and a test set, with blood plasma and serum processed from all donations. (**f**) Individual identification accuracy utilizing the Lasers4Life-LG cohort as a basis for training and testing. Left panel depicts the accuracy of classifiers trained on experimental plasma fingerprints (black) and simulated plasma fingerprints (blue) - testing both classifiers on experimental plasma fingerprints. Right panel depicts the accuracy of a classifier trained on experimental plasma fingerprints (black) and simulated fingerprints of plasma/serum mixtures (blue) - testing both classifiers on experimental serum fingerprints.

The IR spectra of plasma and serum share many characteristics, due to their relatively similar molecular profiles (Fig. 4**b**). The main variations stemmed from the plasma preparation process, which involved the use of ethylenediaminetetraacetic acid (EDTA), as demonstrated in previous work [29]. To achieve effective classification flexibility between specimens, their differences must be well-characterized. This can be achieved by calculating differences between experimental plasma and serum measurements of the same collected blood sample (Fig. 4**c**). Following the CODI framework, we incorporated such characterized differences into an independent experimental dataset of plasma spectra to generate simulated spectra that resemble a mixture between the specimens (Fig. 4**d**).

Next, we revisited the task of identifying individuals as a readout metric to estimate the capacity of cross-specimen generalization. We employed the Lasers4Life-LG cohort (Fig. 4**e**), where both serum and plasma were available from the same individuals at all blood sampling donations. The dataset was split into a training set, consisting of 4-12 donations per individual, and a test set, consisting of the remaining follow-up donations. As a benchmark, we first examined the efficacy of training two classifiers - one trained on experimental plasma spectra and one trained on simulated plasma spectra, testing both on plasma spectra (Fig. 4**f**, left panel). This investigation confirmed the above-mentioned discovery - in that, with sufficiently large experimental training sets, both experimental-based and simulation-based classifiers perform similarly.

We then applied the same classification procedure as above, but here testing on serum measurements (Fig. 4**f**, right panel). For this analysis, we employed the CODI framework to generate a training set of plasma/serum mixtures. Remarkably, we found that the simulation-based approach had a significant advantage over the experimental approach. A substantial drop in prediction efficiency was observed for the classifier trained on experimental plasma, achieving an accuracy of 0.34. The classifier trained on simulated mixture data nearly fully recovered the initial classification efficiency – achieving an accuracy of 0.89. This unexpected finding demonstrated that the CODI framework enabled the creation of a dataset that can even be robust to variations in biological specimen characteristics.

Altogether, this proof-of-principle analysis further demonstrated the CODI framework’s potential to overcome conventional limitations in biological and biomedical research, enabling new applications. For one, there is no need to re-collect a large number of specimens when deviations occur in the sample collection procedure. One can leverage a limited set of measurements that characterize differences between specimens, collected in a class- and task-independent fashion. The CODI framework may then be employed to extend ML applications to different specimen variations. Nevertheless, here we only demonstrated such potential on serum and plasma. If the specimens widely vary in their molecular composition and reflection of physiology (e.g., blood-based vs. urine- or saliva-based media), this approach may not perform as effectively as demonstrated here. A promising avenue for future exploration may involve adapting a classifier trained on EDTA plasma for use with citrate samples [39].

### Generalization to independently acquired datasets

A crucial aspect in determining how well a medical diagnostic assay is likely to perform is to test it on unseen samples. In biodiagnostic applications, a cross-validation procedure is commonly applied to get an estimate of true (external) classification performance. However, if a bias exists in the collected dataset, e.g., confounding information caused by measurement “batch effects”, the estimated performance may not be reproduced when the classifier is truly externally validated [40]. To test this in a relevant medical application, we considered our previous work [30] - in which four cancer entities were classified against non-symptomatic cancer-free controls. In contrast to our prior work which employed cross-validations [30], here the samples were initially split into a training and a test set that were measured independently (Fig. 5**a**, left panel).

**Figure 5.**
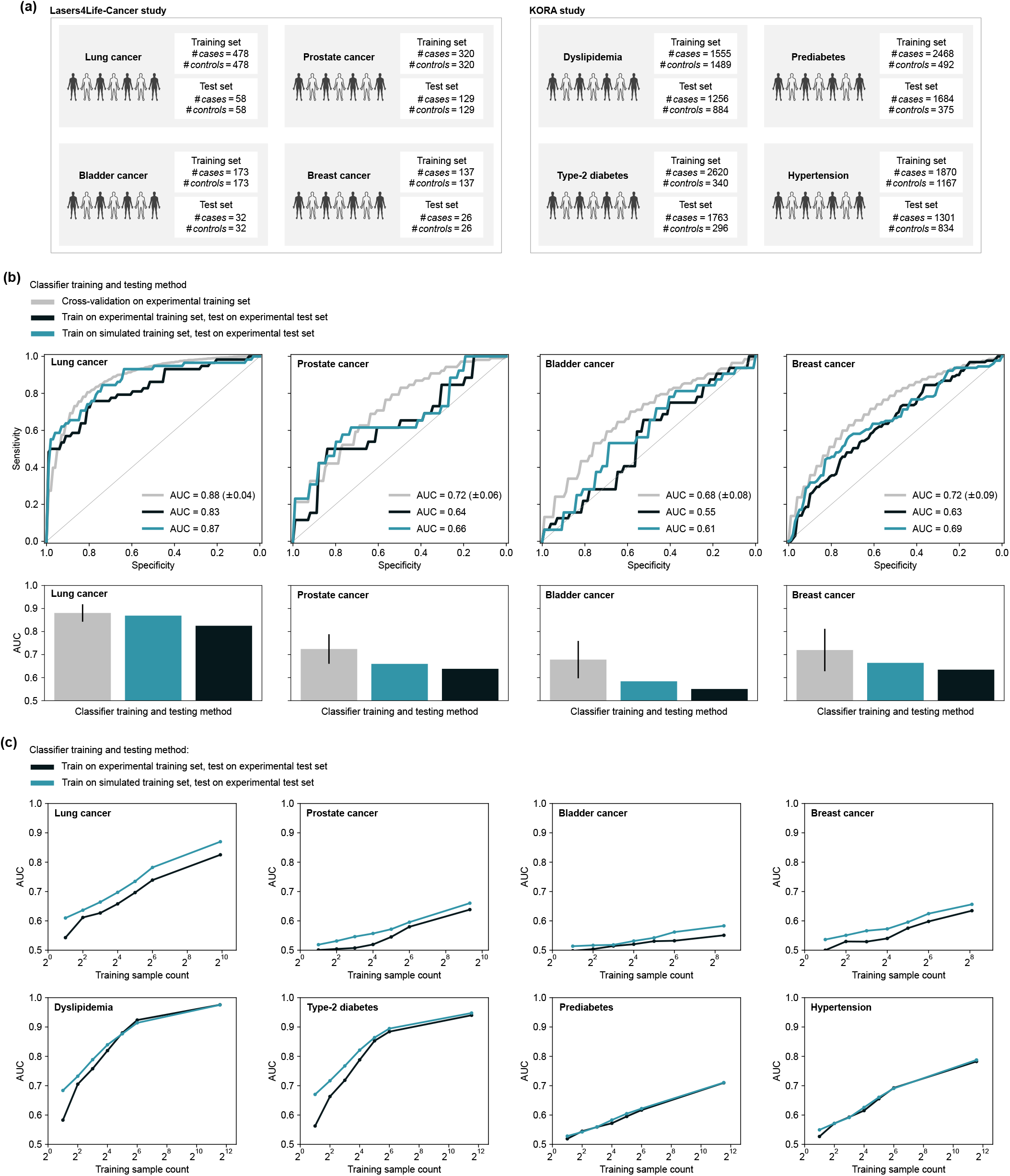
CODI recovers lost classification efficacy on independent case-control test sets. (**a**) Setup of eight binary classifications, spanning diverse health conditions, in two independent clinical studies. IR spectroscopy of blood plasma was performed on all samples. Training and test sample sets were measured in different measurement campaigns under different measurement device conditions. (**b**) Cancer detection was investigated under three different setups of estimating classification efficiency. For the simulation-based training, the CODI framework was employed to introduce measurement variability into the training set measurements. ROC curves are depicted for the validation splits of the data (upper panel), along with the estimated AUCs (lower panel) for each cancer detection application. (**c**) Classification efficiency when training a classifier on experimental observations (black curves) and applying the CODI framework to train a classifier on simulated data (blue curves) utilizing varying training sample counts as a basis for training. Classifier testing was performed exclusively on held-out experimental observations.

As a benchmark, we first investigated the performance of classifying each cancer entity, relying on a cross-validation procedure (Fig. 5**b**, gray curves/bars). The cross-validation was performed exclusively on the training set of experimental samples and the receiver operating characteristics (ROC) curve of the validation splits was examined for each cancer entity. Next, we trained a classifier on the training set of experimental samples, testing it on the test set of experimental samples (Fig. 5**b**, black curves/bars). This investigation revealed that the classification efficiency across the four cancer entities decreased when tested on the later-measured samples. This validated the prior notion that the cross-validation estimates could not be entirely reproduced with this train-test split classification setup.

Next, we questioned whether the CODI framework can aid in such a scenario. In principle, by introducing class-invariant empirical variability into a training set of measurements, we can practically make the learning task more difficult for the classifier. Potentially, this would enable the classifier to appropriately weigh features that are more robust to measurement artifacts, making it rely on information that is likely to be reproduced in unseen data.

To test this, we employed the CODI framework to introduce an additional level of variance to the training set of measurements (Supplementary Information). Across the four cancer entities, we revealed that an improvement in prediction efficiency was indeed observed when testing the classifiers on the held-out test sets (Fig. 5**b**, blue curves/bars). The most impressive improvement was for the lung cancer application, where the area under the ROC curve (AUC) was nearly fully recovered and was comparable to the prior cross-validation estimate. For the remaining cancer entities, the CODI framework still provided an advantage, though not to the same extent as with lung cancer. This may be partly attributed to the occurrence of measurement artifacts in the training set that happened to correlate with the outcome of interest, leading to an overly optimistic AUC estimate during cross-validation. It may also be, in part, due to the generally smaller sample sizes used for testing the classifier and the randomly selected test samples included cases and controls that were more difficult to distinguish than those in the training set (e.g., due to inherent physiological variations that interfere with the cancer signals).

Nevertheless, compared to directly training on experimental observations, the CODI framework consistently led to improved classification output on independently measured test sample data.

### Influence of experimental training cohort size

To investigate under what conditions the CODI framework enables more robust classifier training, we repeated the above case-control investigations, but varied the number of experimental observations utilized for training (Fig. 5**c**). In addition to the previous cancer applications, here we also involved case-control applications from the KORA cohort [41] - focusing on detecting common health physiologies (Fig. 5**a**, right).

We randomly selected samples at varying cohort sizes to train several classifiers on each subset of selected samples (Supplementary Information). First, we trained directly on the experimental observations, always testing on the held-out test sets (Fig. 5**c**, black curves). Unsurprisingly, the smaller the training set was, the worse the classifier performed on the experimental test sets. We then employed the CODI framework to generate simulated datasets that utilized the experimental observations at each sample count as seed input (Fig. 5**c**, blue curves). Here, it was revealed that CODI almost consistently outperformed the experimental modeling approach across the varying sample counts available as a basis for training. Notably, for the detection of dyslipidemia and type-2 diabetes, two conditions with strong molecular deviations reflected in IR fingerprints, CODI provided the largest advance when smaller training sample counts were available.

For the detection of prediabetes and hypertension, no clear advantage was observed by incorporating CODI into the classification workflow. This could either stem from the experimental training data already closely resembling the test data distribution, or because the variability introduced by CODI failed to effectively capture the distribution shifts present in the test data. While no advantage was observed for these two conditions, including CODI did not have an adverse impact on the classification. This observation suggested that integrating CODI into the classification pipeline may be an effective standard practice – as it either enhances prediction performance or, at minimum, does not impair it.

Altogether, our findings show exciting promise for the proposed CODI framework. The value of the method has been demonstrated in the context of several biomedically relevant applications, where the method achieved improved ML classification output for several practical applications.

## DISCUSSION

Multimolecular profiling and computational modeling offer promising avenues to advance our understanding of biological systems. In this study, we introduced CODI, a modeling framework designed to enrich collected datasets to facilitate robust analytics for enhanced levels of system probing. Across several experimental settings, we rigorously tried and tested the framework within the context of molecular profiling to demonstrate its validity. We examined how different empirical and biological factors led to variations in data measured through IR spectroscopy, and revealed the framework’s advantage in overcoming the limitations of unrepresentative observational datasets.

Effectively, the datasets generated through CODI enable an ML algorithm to better capture latent information that is present in a studied dataset, guiding it to distinguish what features are most relevant and reproducible. Such a strategy is particularly valuable when collecting large, representative datasets presents a limitation. This is exemplified in the context of studying pathophysiological phenomena through molecular profiling (e.g., omics studies). In such instances, biological experimentation or medical studies demand substantial involvement, including the probing of a significant number of subjects, considerable time for phenotypic evolution, and the added constraints of intricate sample collection and handling [1–4, 42–44]. Another layer of complexity comes from the fact that biological variations at an organismal level are inherent and inevitable - due to the dynamics of biological systems and human physiologies (e.g., recycling, turnover, rhythmic oscillations, aging) [4, 45–49]. These challenges are further compounded by the often involved quantitative measurement procedures. Factors like the wear and tear of measurement device components, routine maintenance, and sensitivity to environmental conditions all may lead to known “batch effects” that are often specific to a quantitative analytical approach [5, 6, 15, 50, 51]. Ultimately, these challenges impede the generalization of insights to unseen, later collected and measured samples.

The concept of CODI to facilitate generalization draws on data augmentation techniques, which rely on modifying existing experimental observations to create synthetic data. Such techniques find widespread applications in image classification, where data augmentation includes geometric transformations (e.g., rotation, skewing, cropping), color adjustments, and introducing random noise into a training set of images to capture potential variations in practical applications [25]. Data augmentation has also been applied to measurements of biological signals from electroencephalography (EEG), electromyography (EMG), Raman, and near-IR spectra [52–59]. While existing methods often involve random noise introductions, signal warping, and decomposition of available datasets, our approach extends the concept of data augmentation. We take advantage of additional, independent measurements that were not initially a part of the original dataset of interest to model *real* empirical variations for the involved processes of molecular analytics.

In our investigations, we addressed the topic of barely supervised learning [60] - where the set of labeled training samples is limited to very few observations per class. Given the experimental constraints of populational sampling over year-long time-frames, we examined whether the number of sequential samplings of the same individual over time could be minimized with CODI. With the example of IR fingerprinting, we surprisingly identified that only a single baseline measurement is sufficient to follow-up and identify an individual in a population at a later time point. Although identification of the same individual in a heterogeneous population is only a distant approximation to identifying physiologically-relevant deviations, it presents a foundational addition to the concept of longitudinal probing. Our generic framework can be quickly adopted to possibly spare unnecessary samplings and thus inform future prospective studies.

Further applications of CODI to IR spectroscopic fingerprinting showcased its versatility and potential impact in aiding model generalization. We observed a remarkable level of comparability across experimental data collected over almost a decade, underlining the method’s ability to boost classification efficacy on independently measured test sets. The adaptability of the framework extended to a proof-of-principle application that involved training a classifier on one sample medium (plasma) and applying it to another (serum). This application demonstrated the potential of the CODI framework to streamline cross-specimen dataset analyses in various biological and biomedical applications. Such a strategy may be particularly valuable when gathering data sets from retrospective studies or online repositories to help ensure specimen comparability to another envisioned application - i.e., domain adaptation applications [61, 62]. Another promising use of CODI may be to harmonize data obtained from different measurement devices (e.g., spectrometers made by different manufacturers), potentially improving ML model transferability between them [9].

It is imperative to emphasize that our proposed method shall not be positioned as a replacement for improving study designs and better standardization of classical analytical procedures. Such aspects remain essential when establishing a molecular profiling platform to satisfy i.i.d. assumptions between train-test datasets. In addition to efforts aimed at ensuring train-test dataset comparability, the motivation of our method is to work around the instances when the assumption is violated, due to inevitable sources of error and variability.

An inherent limitation of the proposed method is its reliance on *a priori* knowledge about the sources of possible empirical variations. This presents a challenge as gathering such information may require extensive experimental evaluations of biological and analytical variations. Furthermore, the characterized sources of variability must adequately represent the true possible domain of empirical variability for any successful application. Therefore, continuous refinement through more controlled experiments to include several, independent, sources of variance holds potential to further enhance the utility of the framework. Nevertheless, once the domain of possible variance is successfully characterized for a given system and measurement procedure, the same characterizations may be repeatedly utilized in diverse applications.

While our investigations primarily focused on blood-based IR spectroscopy to aid *in vitro* diagnostics, the practical applications of the CODI framework are not limited to this context. The principle and mathematical foundation of the framework is sufficiently generic to be translated to the examination of diverse biological systems, medical problems, measurement modalities, and ML tasks. Applications involving NMR spectroscopy, mass spectrometry, and Raman spectroscopy serve as direct extensions that can be explored with this modeling framework. Additionally, the framework holds potential in applications related to cell typing and the integration of single-cell multimodal omics data – given the inherent challenges associated with obtaining accurate measurements of cell type populations at scale, where out-of-distribution measurement events are prevalent [63, 64]. Altogether, the presented framework establishes a foundation for future explorations to enhance the robustness of molecular analytics.

## MATERIALS AND METHODS

More detailed descriptions of the CODI modeling framework, our applications, datasets, experimental procedures, and ML analysis are provided in the Supplementary Information.

In brief, the *in silico* model behind CODI is designed for extensibility, such that it can be applied in various applications and for different measurement modalities. The modeling framework involves utilizing seed observations {***s***_*i*_ | *i* = 1, …, *m*} that are representative of properties intrinsic to a phenomena of interest. For instance, a seed observation might represent the mean observation of a class of samples (e.g., healthy control), while another corresponds to a different class (e.g., disease sample).

Variability is introduced to the seed observations ***s***_*i*_ through the addition of random functions *f*_1_, *f*_2_, …, *f*_*m*_, simulating a measurement as a statistical variable ***Y*** in the following generalized form:

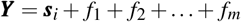

Repeatedly applying the above model would therefore generate a cohort of simulated measurements, in arbitrary size, centered around ***s***_*i*_ and incorporating the variations introduced by *f*_1_, *f*_2_, …, *f*_*m*_.

The variations introduced by *f*_1_, *f*_2_, …, *f*_*m*_ can be characterized by *ab initio* calculations or bottom-up models that each represent a source of expected data variance. However, the former is often specialized and problem-specific. An alternative descriptive approach that relies on collecting datasets of calibration measurements which incorporate the levels of expected variance can be easily applied to a variety of problems. For instance, quality control samples can be subjected to different freezer-storage durations, number of freeze/thaw cycles, and aliquoting of samples by different operators. The quality control samples can then be repeatedly measured under different measurement device conditions. The variance observed in this calibration dataset would be reflective of potential sources of variance in handling and measuring samples from the original dataset modeling a certain phenomena (e.g., biofluids of cases and controls). This variance can then be introduced by defining the function *f*_1_. Other potential sources of data variance, such as biological variability, can be modeled by using additional calibration datasets reflective of the variability sources.

Four independent study cohorts were utilized in our applications of CODI: Lasers4Life-LG [29], BioPersMed [27], KORA [28, 41], and Lasers4Life-Cancer [30]. Lasers4Life-LG involved the collection of blood serum and plasma from 31 healthy individuals initially sampled up to 13 times over a 7-week period, with an additional follow-up after 6 months as detailed in a previous publication [29]. Since this initial publication, the same individuals were invited to participate in two additional sample donations at 3.5 and 4.5 years post their initial involvement. BioPersMed is an ongoing population-based cohort performed at the Medical University Graz, Austria [27]. Repetitive examinations of participants were conducted in 2-year intervals. In the current study, we utilized blood plasma samples and medical data from a subset of 44 healthy individuals (out of 1022 participants). KORA is a population-based cohort in Southern Germany [28]. The cohort comprised of an age- and sex-stratified sample of participants randomly drawn from the resident registration offices within the study area. In the current study, we utilized blood plasma samples and medical data from the second and third visits (named KORA-F4 and KORA-FF4, respectively). The available KORA-F4 data consisted of 3044 samples, while the KORA-FF4 data consisted of 2140 samples. A subset of 2015 individuals participated in both samplings, while 1154 individuals participated in only one of the samplings. Lasers4Life-Cancer is a case-control study cohort involving several cancer entities were both serum and plasma are collected [30]. The samples utilized in this study largely overlapped with samples from our previous study that involved the detection of four common cancer entities (lung, prostate, bladder, and breast) [30] – although the measurement procedures differed (Supplementary Information). Since the original publication, blood plasma and serum samples from different individuals were collected and included in this study. Case samples were collected prior to cancer-related treatment (i.e., therapy-naive). Non-symptomatic controls were pair-matched to cancer cases by age, sex, and body mass index.

Experimental measurements of the liquid samples were performed on a Fourier transform IR (FTIR) spectrometer as in previous studies [29, 30, 41]. Samples were injected into a flow cell, and the IR spectra were recorded in transmission mode. The measurement device underwent routine maintenance, with certain components replaced as needed throughout measurements of all clinical samples (Supplementary Information).

Multi-class classifications for individual identification were performed using a nearest neighbor algorithm for applications involving training with a single observation per class. For multiclass applications involving training more than one observation per class, a linear discriminant analysis (LDA) algorithm was applied. Binary classifications for case-control applications were performed using a logistic regression algorithm with a ridge penalty. All metrics of classification efficacy were reported on held-out test experimental samples (Supplementary Information).

## CODE AVAILABILITY

An implementation of CODI is available as a Python package. Please refer to the following GitHub repository for details on its usage: https://github.com/tarek-eissa/codi.

## AUTHOR CONTRIBUTIONS

T.E. conceived the project; T.E., M.H., and M.Ž. contributed to study design; T.E. performed the research and analyzed the data with assistance from M.H.; M.Ž ., B.O.P., B.L., and A.P. oversaw, organized, and coordinated the clinical studies; F.F. supervised and contributed to the experimental measurements and coordinated data transfers; T.E. and M.Ž. wrote the manuscript with assistance from M.H. and F.F.; All authors revised the manuscript. All authors approved the manuscript.

## ACKNOWLEDGMENTS

The KORA study was initiated and financed by the Helmholtz Zentrum München – German Research Center for Environmental Health, which is funded by the German Federal Ministry of Education and Research (BMBF) and by the State of Bavaria. Data collection in the KORA study is done in cooperation with the University Hospital of Augsburg. The BioPersMed cohort study was funded by the Austrian Research Fund, as a COMET K-project (No. 825329), supported by the Austrian Federal Ministry of Transport, Innovation and Technology (BMVIT) and the Austrian Federal Ministry of Economics and Labour/the Federal Ministry of Economy, Family and Youth (BMWA/BMWFJ) as well as the Styrian Business Promotion Agency (SFG). We are grateful to Ferenc Krausz for the research environment, Jacqueline Hermann for clinical study management and for reviewing the manuscript, Viola Zoka and Daniel Meyer for technical contributions, and Michael Trubetskov for reviewing the manuscript.

—–

The publication and the software are subject to copyright and belong to the Max Planck Society and Ludwig Maximilian University of Munich. The content of the publication is subject of a patent application.

