## Supplementary Information for "CODI: Enhancing machine learning-based molecular profiling through contextual out-of-distribution integration"

### 1 SUPPLEMENTARY METHODS

#### SIMULATION MODEL

In the following subsections, we provide a general framework of the simulation model behind CODI to facilitate its extensibility to other applications and molecular fingerprinting modalities. Technical details on our specific applications of the model are provided later in the text.

##### Generalized representation

CODI relies on introducing information about possible sources of variations (biological, technical, etc.) that may arise in an experimental setting to an existing set of experimental observations $\{\boldsymbol{x}_i \mid i = 1, \ldots, n\}$ where each $\boldsymbol{x}_i$ is a numerical vector. In a generalized form, the sources of variability can be denoted as functions $f_1, f_2, \ldots, f_m$, each representing a model for distinct aspects of variation. These functions are assumed to be random vectors in the same space as the experimental observations, and taking a probability distribution centered around 0.

From the set of experimental observations $\boldsymbol{x}_i$, we can extract another set of experimentally-derived seed observations $\{\boldsymbol{s}_i \mid i = 1, \ldots, m\}$ that model intrinsic properties of a sub-group in the original set of data – e.g., different classes. With these, a resulting simulated measurement can be modeled as a statistical variable $\boldsymbol{Y}$, using the following generalized form:

$$\boldsymbol{Y} = \boldsymbol{s}_i + f_1 + f_2 + \ldots + f_m \tag{1}$$

Repeatedly applying the above model would therefore generate a cohort of simulated measurements, in arbitrary size, centered around $\boldsymbol{s}_i$ and incorporating variations introduced by $f_1, f_2, \ldots, f_m$.

##### Setting the seed measurements

Defining the experimentally-derived seed measurements $\boldsymbol{s}_i$ depends on the available input dataset of experimental measurements $\boldsymbol{x}_i$ and the type of measurement event we wish to simulate.

In a straightforward implementation, we can set $\boldsymbol{s}_i = \boldsymbol{x}_i$. This formulation would allow us to introduce a level of variability to each measurement-specific observation. Repeatedly applying the model for each $\boldsymbol{x}_i$ would create several sets of measurements, where each set is centered around each experimental observation. In an alternative definition, we can set $\boldsymbol{s}_i = \bar{\boldsymbol{x}}$. This would allow us to introduce a level of variability to the mean measurement of the experimental set $\boldsymbol{x}_i$, thereby, creating a cohort of measurements centered around their expected value.

If all measurements in $\boldsymbol{x}_i$ reflect a certain group or class of samples (e.g., healthy individuals), the simulated cohort as a whole would reflect a measurement distribution of that class of samples. Trivially, when the dataset $\boldsymbol{x}_i$ is switched to involve measurements representative of a different class (e.g., disease cases), the simulated cohort would reflect a distribution centered around the alternative outcome.

##### Introduction of variability modeling functions

The variations introduced by the functions $f_1, f_2, \ldots, f_m$ may be characterized by *ab initio* calculations or bottom-up models that each represent a source of expected data variance. However, the former is highly specialized and problem-specific. An alternative descriptive approach that relies on collecting datasets of calibration measurements which incorporate the levels of expected variance can be easily applied to a variety of problems.

Suppose an independent calibration dataset $\{\boldsymbol{b}_i \mid i = 1, \ldots, l\}$ is available that reflects a given source of measurement variability. Here, $f_1$ would take the following form:

$$f_1 = \sum_{i=1}^{l} \beta\left(0, \frac{1}{\sqrt{l}}\right) \cdot (\boldsymbol{b}_i - \bar{\boldsymbol{b}}) \tag{2}$$

By scaling and combining $l$ individual deviations from the mean measurement $(\boldsymbol{b}_i - \bar{\boldsymbol{b}})$ using a Gaussian random variable $\beta\left(0, \frac{1}{\sqrt{l}}\right)$ centered around 0, the random function $f_1$ would have the same variance as the dataset $\boldsymbol{b}_i$. The formal and empirical proof for this was described in a previous study where $f_1$ generated a level of measurement variance that is observed between different individuals [1]. If the variance source in question does not follow a Gaussian distribution, the random variable $\beta$ may assume a more appropriate probability distribution.

Additional functions $f_2, f_3, \ldots, f_m$ may be modeled similarly by utilizing different calibration datasets that reflect other sources of measurement variability. In our example applications of the model in this study, the utilized calibration datasets reflected levels of within- or between-person biological variability, along with several sources of analytical variability.

Descriptions on the model variants applied in our applications of the model, along with of how the calibration datasets were defined, is detailed in following sections of the text.

##### Simulating case-control measurements

For analysis that involved simulating measurements of cases and controls in our applications (e.g., detection of cancer), the input dataset took the following form, where the superscript denotes a case or control:

$$\boldsymbol{D} = \left\{\boldsymbol{x}_1^{(+1)}, \boldsymbol{x}_2^{(-1)}, \boldsymbol{x}_3^{(-1)}, \ldots, \boldsymbol{x}_n^{(+1)}\right\} \tag{3}$$

Two model definitions were applied based on Eq. 1, once by setting $\boldsymbol{s}_i = \bar{\boldsymbol{x}}^{(+1)}$ and once by setting $\boldsymbol{s}_i = \bar{\boldsymbol{x}}^{(-1)}$. This results in the following model formulations:

$$\begin{aligned} \boldsymbol{Y}^{(+1)} &= \bar{\boldsymbol{x}}^{(+1)} + f_1 + f_2 + \ldots + f_m \\ \boldsymbol{Y}^{(-1)} &= \bar{\boldsymbol{x}}^{(-1)} + f_1 + f_2 + \ldots + f_m \end{aligned} \tag{4}$$

Both model variants were then repeatedly applied to generate a dataset containing case and control measurements for each disease detection application separately.

#### Simulating longitudinally-captured measurements

For an application that involved simulating longitudinal measurements of the same individuals, the input dataset took the following form:

$$\boldsymbol{D} = \left\{ \boldsymbol{x}_0^{(a)}, \boldsymbol{x}_1^{(a)}, \boldsymbol{x}_2^{(a)}, \boldsymbol{x}_0^{(b)}, \ldots, \boldsymbol{x}_0^{(z)}, \boldsymbol{x}_1^{(z)} \right\} \tag{5}$$

where the superscript denotes different individuals and the subscript denotes their visit number.

For each individual, we may then utilize their baseline measurement (visit 0) to simulate a set of measurements centered around that baseline in the following form:

$$\begin{aligned} \boldsymbol{Y}^{(a)} &= \boldsymbol{x}_0^{(a)} + f_1 + f_2 + \ldots + f_m \\ \boldsymbol{Y}^{(b)} &= \boldsymbol{x}_0^{(b)} + f_1 + f_2 + \ldots + f_m \\ &\vdots \\ \boldsymbol{Y}^{(z)} &= \boldsymbol{x}_0^{(z)} + f_1 + f_2 + \ldots + f_m \end{aligned} \tag{6}$$

Rather than only using the baseline measurements, the same concept can be extended to incorporate additional measurements obtained during subsequent visits for each individual. The process can therefore generate an arbitrary number of measurements for each individual – whether based on a single baseline measurement or more.

#### Simulating between-person biological variability

To model variations in measurements between different individuals, the calibration dataset $\boldsymbol{b}_i$ should include experimentally obtained measurements of different individuals. Ideally, all measurements captured for different individuals should be performed in a similar experimental workflow, following the same analytical protocol. This helps ensure that the variance of the calibration dataset is driven primarily by the biological differences between different individuals, excluding the contributions of other sources of data variance. Here, $f_1$ may be set to model the between-person biological variability and would be defined in the same way described in Eq. 2.

#### Simulating within-person biological variability

To model variations in measurements of the same individual over time, the calibration dataset $\boldsymbol{b}_i$ should include several experimentally obtained measurements per individual. Similar to the longitudinal dataset definition provided in Eq. 4, we can formulate another calibration dataset of $l$ entries that model within-person deviations in the following form:

$$\boldsymbol{D} = \left\{ \boldsymbol{b}_0^a - \bar{\boldsymbol{b}}^a, \boldsymbol{b}_1^a - \bar{\boldsymbol{b}}^a, \boldsymbol{b}_2^a - \bar{\boldsymbol{b}}^a, \ldots, \boldsymbol{b}_0^z - \bar{\boldsymbol{b}}^z, \boldsymbol{b}_1^z - \bar{\boldsymbol{b}}^z \right\} \tag{7}$$

In other terms, the calibration vectors were calculated per individual and the whole set of individual-specific deviations can be pooled into one dataset.

Utilizing the dataset defined in Eq. 7, the function $f_1$ may be set to model within-person biological variability and would be defined similarly to Eq. 2 with the following form:

$$f_1 = \sum_{\boldsymbol{d} \epsilon \boldsymbol{D}} \beta \left( 0, \frac{1}{\sqrt{l}} \right) \cdot \boldsymbol{d} \tag{8}$$

With this, the function $f_1$ thus models the within-person variability as observed across multiple individuals.

#### Simulating analytical variability

To model variations that may arise from a given analytical procedure, additional experimental measurements that are subjected to experimental variations introduced during the analytical procedure can be utilized. This includes variations that may arise from sample preparation, environmental factors, and instrumental errors. In general, describing the reproducibility of collected data can be achieved by conducting multiple measurements of replicates. Specifically defining how to handle and measure the replicates is dependent on the analytical workflow applied and the measurement technique.

Measurements of replicates can be performed on samples with a high level of technical reproducibility (e.g., water samples) or on quality control samples that exhibit stability in chemical composition similar to that of the intended application (e.g., blood-based samples pooled from thousands of individuals). With blood-based quality control samples, for instance, variations could further include freezer-storage durations, number of freeze/thaw cycles, aliquoting of samples by different operators, varied experimental laboratory room temperature, and sample storage in tubes from different manufacturers.

Technicalities specific to the measurement device may also be intentionally varied when performing measurements – e.g., filter changes, software versions, and cuvette lifetimes. The replicate samples may also be performed across multiple measurement devices, and environmental conditions, including temperature and humidity, to further enrich a calibration dataset.

The variations that arise from the repeated replicate measurements would then be modeled in the same fashion described in Eq. 2. $f_2$, for instance, may encompass all the sources of measurement variability that are reflected in a blood-based quality control sample, while, $f_3$ may encompass the sources of measurement variability that are reflected in water measurements.

Modeling such factors, in a way that best mimics what may be encountered in application, makes a given training dataset more resilient to molecular changes or measurement device operating conditions that may arise from an analytical workflow.

#### Defining which variability functions to include

Successful applications of the CODI modeling framework relies on the careful selection of appropriate sources of variability $f_1, f_2, \dots, f_m$. This selection depends on the application of interest and what sources of measurement variability are expected to be observed. For instance, if the envisioned application is to model case-control measurements, it is more critical to include a level of between-person biological variability than to include a level of within-person biological variability. Conversely, if the goal is to simulate how an individual-specific measurement varies over time, including a level of within-person biological variability is necessary. Sources of analytical errors may be independent of the envisioned application of an analytical procedure (e.g., measurement device drifts, or sample storage variations). Therefore, such variability functions may be included, regardless of the application setting.

In a model validation phase, where experimental test samples are available, it must be ensured that the calibration datasets utilized do not include any of the test samples. This avoids leaking any information or statistical properties of the test samples into the training samples.

In our applications, the variations between quality control samples, between different studies, and between technical replicate measurements were always included in the model. The within-person biological variability was only included for longitudinal applications, while the between-person biological variability was included for case-control applications. The variations between plasma and serum were only included when transferring a model trained on plasma measurements to its application on serum measurements. Descriptions of what calibration measurements were utilized for each application are provided in the following sections of the text.

### CLINICAL STUDIES

#### Lasers4Life-LG study

The Lasers4Life-LG study cohort comprised of 31 nominally healthy individuals, initially sampled up to 13 times over a 7-week period, with an additional follow-up after 6 months as detailed in a previous publication [2]. Since this initial publication, the same individuals were invited to participate in two additional sampling sessions at 3.5 and 4.5 years post their initial involvement. In both the latter two samplings, 8 individuals participated. Blood plasma and serum were collected from all participants throughout the study. The study was approved by the Ethics Committee of the Ludwig-Maximillian-University (LMU) of Munich and all participants provided written informed consent (research study protocol #17-532).

#### BioPersMed study

The BioPersMed study is an ongoing population-based cohort at the Medical University Graz, Austria [3]. Repetitive examinations of participants were conducted in 2-year intervals. In the current study, we utilized blood plasma samples and medical data from a subset of 44 nominally healthy individuals out of 1022 participants. All 44 individuals participated in the baseline sampling, and follow-up visits ranged up to 8 years. Further detailed description of the study design was previously published [3]. The study was approved by the Ethics Committee of the Medical University of Graz, Austria (EC Nr. 24-224 ex 11/12; project application number 4008_22).

#### KORA study

The KORA study is a population-based cohort in Southern Germany [4, 5]. The study comprised of an age- and sex-stratified sample of participants randomly drawn from the resident registration offices within the study area. In the current study, we utilized blood plasma samples and medical data from the second and third participant visits (named KORA-F4 and KORA-FF4, respectively). The available KORA-F4 data consisted of 3044 samples, while the KORA-FF4 data consisted of 2140 samples. A subset of 2015 individuals participated in both samplings, while 1154 individuals participated in only one of the samplings. The samplings were separated by an average of 6.5 follow-up years. Data collection methods and standardized sample collections have been described in detail elsewhere [4, 6–8]. The KORA-F4 and KORA-FF4 study methods were approved by the ethics committee of the Bavarian Chamber of Physicians, Munich (EC No. 06068).

#### Lasers4Life-Cancer study

Samples utilized for the cancer analysis were derived from the case-control Lasers4Life-Cancer study. The samples utilized in this study largely overlap with samples utilized in our previous study that involved the detection of four common cancer entities (lung, prostate, bladder, and breast) [9]. Newly collected blood plasma and serum samples from different individuals were included to increase the sample size and thus improve the statistical robustness of the results. Case samples were collected prior to any cancer-related treatment (i.e., therapy-naive). Non-symptomatic individuals were used as controls. For each cancer entity, controls were pair-matched to the cancer cases by age, sex, and body mass index (BMI). All participants provided written informed consent for the study under research study protocol #17-141 and under research study protocol #17-182, both of which were approved by the Ethics Committee of the LMU of Munich. The clinical trial is registered at the German Clinical Trials Register (ID DRKS00013217).

#### EXPERIMENTAL PROCEDURES

##### Infrared spectroscopy

Infrared spectroscopy was performed on liquid samples using Fourier transform IR (FTIR) spectrometer (MIRA Analyzer, CLADE GmbH, Esslingen, Germany). After a sample was injected into a flow cell (window material calcium fluoride) with approx. 8 µm of optical pathlength, the IR spectrum of the sample was recorded in transmission mode and subsequently a reference spectrum of the transport medium (i.e., calcium fluoride saturated water) was recorded. The spectra were acquired with a resolution of 4 $cm^{-1}$ in a spectral range between 950 $cm^{-1}$ and 3050 $cm^{-1}$. The raw data provided by the MIRA Analyzer is the absorption spectrum of the sample subtracted by the reference spectrum. Preprocessing of the resulting IR spectra was performed as described previously [9].

The instrument was maintained by the manufacturer on a yearly basis. The components that were replaced during the yearly or routine user maintenance with potential effect on the IR spectra are the following: light source (after three years); desiccant cartridges (yearly, or when necessary); flow cell when the optical path length exceeded the limit (typically after several thousands of samples), 2 $\mu$m flow cell pre-filter (typically after 200-300 samples).

##### Sample handling

All clinical samples involved were stored at -80°C after being processed to serum or plasma. The transport to measurement laboratory was performed on dry ice, and further storage was at -80°C. Original samples (0.3 to 1.0 mL) were thawed at 4°C, and centrifuged for 10 min at 2000 g. The supernatant was aliquoted into the measurement tubes (50 - 100 µL per tube) and refrozen until measured.

Purchased pooled human serum was used as quality control (QC) samples. 3 liters (100 mL flasks, BioWest, Nuaillé, France) were ordered and stored in original flasks at -80°C. Prior to use, these were thawed at 4°C, filtered through a 0.45 µm filter, and aliquoted to 50-100 µL aliquots that were kept at -80°C until use. Each sample measurement batch started and ended with a QC serum measurement. In addition, a QC sample was measured after every 5 clinical samples. A measurement batch consisted of 25 to 40 samples. Although the same lot of the QC serum was used, slight variability between the flasks were measured, possibly due to the difference in storage duration and technical handling.

To evaluate the technical variability of the FTIR spectrometer, pure water samples were measured. Although not measured daily, several hundreds of water measurements were performed over the years. Due to the pre-processing of measurement data by the FTIR device (i.e., subtraction of the reference spectrum), the resulting deviations reflect the technical noise of the measurement procedure.

##### Measurements of clinical samples

Samples of the Lasers4Life-LG study were measured in a fully randomized order over all visits but separately for serum and plasma (single flow cell). BioPersMed study samples were measured in a fully randomized order (single flow cell). KORA-F4 and KORA-FF4 study samples were measured in two campaigns separated by an average of 2.7 years between the measurement dates of each sampling. Three flow cells were used throughout both measurement campaigns (KORA-F4: single flow cell; KORA-FF4: two flow cells).

For the Lasers4Life-Cancer study, the whole study was split into a training and a test sample set prior to any measurements. Within these sets, the measurement order was fully randomized. The samples of the training set were measured with three flow cells. A gap of four weeks occurred prior to measuring the test set samples. Within the gap, fully independent measurements were routinely performed. Measurements of the test set were performed on another, fourth, flow cell to simulate potential measurement drifts.

All samples were measured in small batches of 25-40 samples plus QC serum as described above. After each measurement batch, the device was cleaned according to the manufacturer's recommendations. A performance qualification test of the instrument was performed daily before the first sample batch or after the clearance of technical issues.

#### SIMULATION AND CLASSIFICATION ANALYSIS

##### Individual identification in longitudinal monitoring

Simulated datasets of longitudinal measurements were generated using the model formulation presented in Eq. 6. For the analyses illustrated in Fig. 3**b–d**, the baseline measurement of each individual served as the initial seed input. In the analysis depicted in Fig. 3**e**, the model formulation in Eq. 6 was iteratively applied, utilizing up to the first 6 baselines per individual as seed inputs.

Four variability functions were employed to introduce variability to each seed input: (1) within-person biological variability; (2) clinical sampling variability; (3) quality control variability; and (4) technical replicate variability. The within-person variability was always modeled by only using data of two of the three utilized longitudinal cohorts (depicted in Fig. 3**a**). When the individual identification classification was investigated in one of the three cohorts, the within-person variability was characterized from the two other cohorts. Since the within-person biological variability relied on utilizing follow-up measurements (Eq. 8), this step ensured that no information was leaked to the training sets from the test sets (the follow-ups). For each experimental baseline, 1000 simulated measurements were generated to ensure a sufficiently large sample size, reaching a plateau of classification performance.

Before classification, a data standardization step (mean is 0, standard deviation is 1) and principle component analysis (PCA) were applied. For the data standardization, the mean and standard deviation were calculated only from the training sets to standardize both the training and test sets. PCA was applied to the training sets - keeping components that explain 99.99% of the total variance. The loading vectors from PCA were then applied to both the training sets and test sets before training. After the preprocessing prior to classification, a linear discriminant analysis (LDA) algorithm was used for multi-class classifications when more than one observation per class was available for training. When only one instance per class was available in the training set, a K-nearest neighbor (KNN) algorithm was for classifications (with $k = 1$). Classifier testing was always performed on held-out samples from follow-ups that were not used to create training datasets. Confidence intervals (CIs) were constructed by bootstrap resampling of the test data, following the approach applied in previous work [10]. The [2.5, 97.5] percentile boundaries were chosen to construct a 95% CI interval of the classification accuracy.

##### Generalization between plasma and serum specimens

For the application that involved creating simulated spectra of plasma and serum mixtures (Fig. 4), a similar simulation and classification workflow was applied as described in the prior section. However, in addition to the four previously described sources of variability, an additional function was introduced to model deviations between serum and plasma measurements. The calibration datasets used to characterize the differences between IR spectral measurements of blood plasma and serum (depicted in Fig. 4**c**) were calculated from the Lasers4Life-Cancer study, where plasma and serum were were collected at the same sampling occasion. The individual identification application was then applied to the Lasers4Life-LG cohort (Fig. 4**f**) – an independent cohort that involved different individuals than the Lasers4Life-Cancer cohort.

##### Classifier testing on independent test sets

An L2-regularized logistic regression algorithm was used for the binary (case-control) classification analysis. Data was initially split into a training and test set (Fig. 5**a**). Three different setups of evaluating the classification efficiency were performed. First, a 10-times repeated 10-fold stratified cross-validation was carried out on the training set of samples. Classification efficiency was evaluated on the validation splits of the cross-validation by calculating the area under the receiver operating characteristic curve (AUC). The AUC was averaged across the validation splits and reported along with its standard deviation. In the second setup, the classifier was trained directly on the experimental data. The AUC was then calculated from the unseen, initially held out, test set. In the third setup, classifier training was performed on a simulated set of samples - based on the training set as seed data. The AUC was again calculated from the unseen, initially held out, experimental test set.

When creating the simulated datasets, four variability functions were employed to introduce variability to each seed input: (1) between-person biological variability; (2) clinical sampling variability; (3) quality control variability; and (4) technical replicate variability. The data sources for calculating the level of between-person variability involved the training set of samples, as well as samples from the other independent studies - i.e., samples from all utilized clinical studies, outside of the test set. The model formulation in Eq. 4 was applied, where the mean measurement of cases and controls served as seed measurements for each binary classification task. For each task, 100000 samples were generated per class.

#### ANALYSIS SOFTWARE

Analyses in this study were performed using custom scripts written in Python (v.3.8.8). The open-source packages NumPy (v.1.21.2), scikit-learn (v.0.24.1), and matplotlib (v.3.5.1) were used. An implementation of CODI is available as a Python package. Please refer to the following GitHub repository for details on its usage: `https://github.com/tarek-eissa/codi`.

#### 2 CHOOSING THE VARIABILITY FUNCTIONS

The sources of variability that we utilized in our applications of CODI were characterized by sets of calibration measurements that reflect a level of data variance. As previously described, within these calibration measurements were characteristics of empirical variability stemming from inherent biological factors, variations in sample collection and handling, as well as instrument-specific measurement noise and drifts. Specifically, in our longitudinal analyses, we characterized four main sources of variability: (1) within-person biological variability, (2) clinical sampling variability, (3) quality control variability, and (4) technical replicate variability. Here, we examined how each of these sources of variability contributed to the success of the classification.

Fig. S1 depicts an analysis where the classification task was to identify an individual given one baseline measurement per individual/class – testing the classification on follow-up measurements. Similar to the analysis depicted in Fig. 3**a**–**b** in the main text, this analysis was also carried out on the three independent longitudinal cohorts. The CODI modeling framework was employed to generate simulated training data based on all four sources of variability (Fig. S1, blue bars). We then systematically removed one of the variability sources considered in the simulation model (keeping the other three) to examine whether the classification accuracy was affected (Fig. S1, remaining bars). We found that with the absence of either the within-person biological variability or the quality control variability, the classification accuracy significantly suffered. Specifically for the application on the samples from KORA clinical study, a very significant drop in the prediction accuracy was observed when the simulation model did not include the variability stemming from the quality control samples. Interestingly, the KORA samples — the baseline measurements and follow-ups – were measured at very different times, years apart (see section "Measurements of clinical samples" above). Given that the quality control measurement include information about potential measurement drifts that may be observed when performing measurements at very different times, this highlighted the importance of the informed inclusion of variance that is characteristic of what may be observed in realistic applications. On the other hand, the variability introduced by the clinical sampling and technical replicate calibration measurements had minimal impact on the classification accuracy – revealing that, for our specific use-cases, these variability sources are not crucial to account for in the modeling process. Nevertheless, no significant loss in the classification accuracy was observed when all four sources of variability were included in the simulation model. Thus, we included all four sources in our analyses.

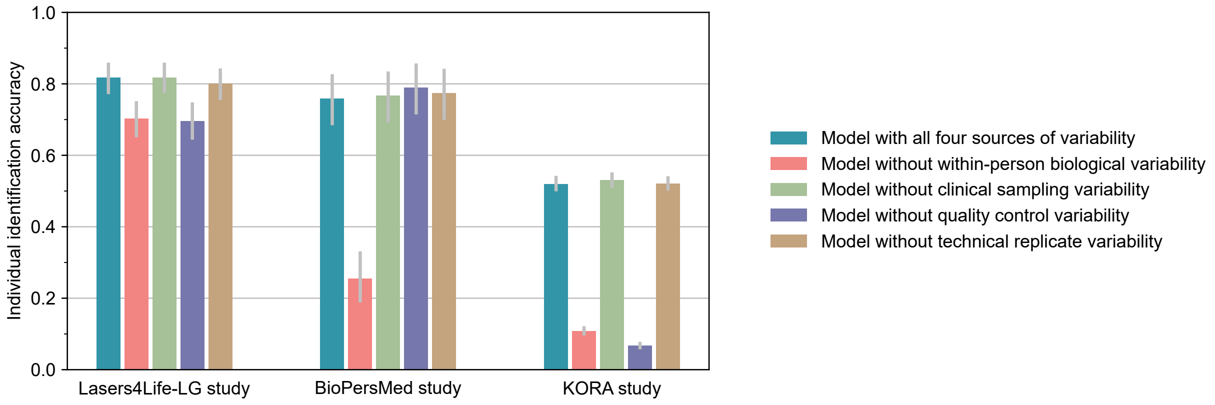


**Figure S1.** Impact of the modeled variability sources on the classification accuracy in longitudinal molecular monitoring. Each bar illustrates the classification accuracy achieved by simulating data through CODI with different combinations of the variability sources included.

#### 3 COMPARATIVE ANALYSIS OF CODI TO DOMAIN-AGNOSTIC AUGMENTATION TECHNIQUES

The CODI modeling framework inherently relies on *a priori* information on possible sources sample/measurement variability. In contrast, domain-agnostic augmentation methods employ generic transformations (e.g., additive or multiplicative noise) on the input data to simulate new measurements, eliminating the need for *a priori* information. Here, we provide a comparative analysis between CODI and other domain-agnostic methods of generating simulated training sets. Specifically, for spectral datasets, domain-agnostic augmentation methods often include the introduction of additive white noise, random scaling of intensities by multiplicative factors, linear slope variations, and vertical offset shifts [11, 12] (Fig. S2**a**).

Fig. S2**b** depicts an analysis where several domain-agnostic augmentation methods were employed to generate simulated training data for the classification task of identifying individuals based on a single baseline measurement per class. As benchmarks, we include the performance of classifiers trained directly on the experimental data and simulated data generated through CODI. Each augmentation strategy, including CODI, was applied to the same (seed) experimental training data, generating 1000 measurements per class, and tested on the same follow-up experimental measurements. Unlike CODI, the domain-agnostic augmentation methods require the tuning of free parameters that control the extent to which they affect the seed input. Across the three longitudinal cohorts this investigation was carried out, we found that the data generated through CODI had the clear advantage over all other methods.

While the domain-agnostic augmentation techniques are simpler to implement and do not require *a priori* knowledge, overall they yielded minimal-to-no improvement in the classification accuracy when compared to training directly on the experimental data. Unlike the domain-agnostic augmentation methods, CODI's targeted integration of unrepresented variations allowed for generating more representative simulated data that enhanced the classifier's ability to generalize beyond the original experimental training set.

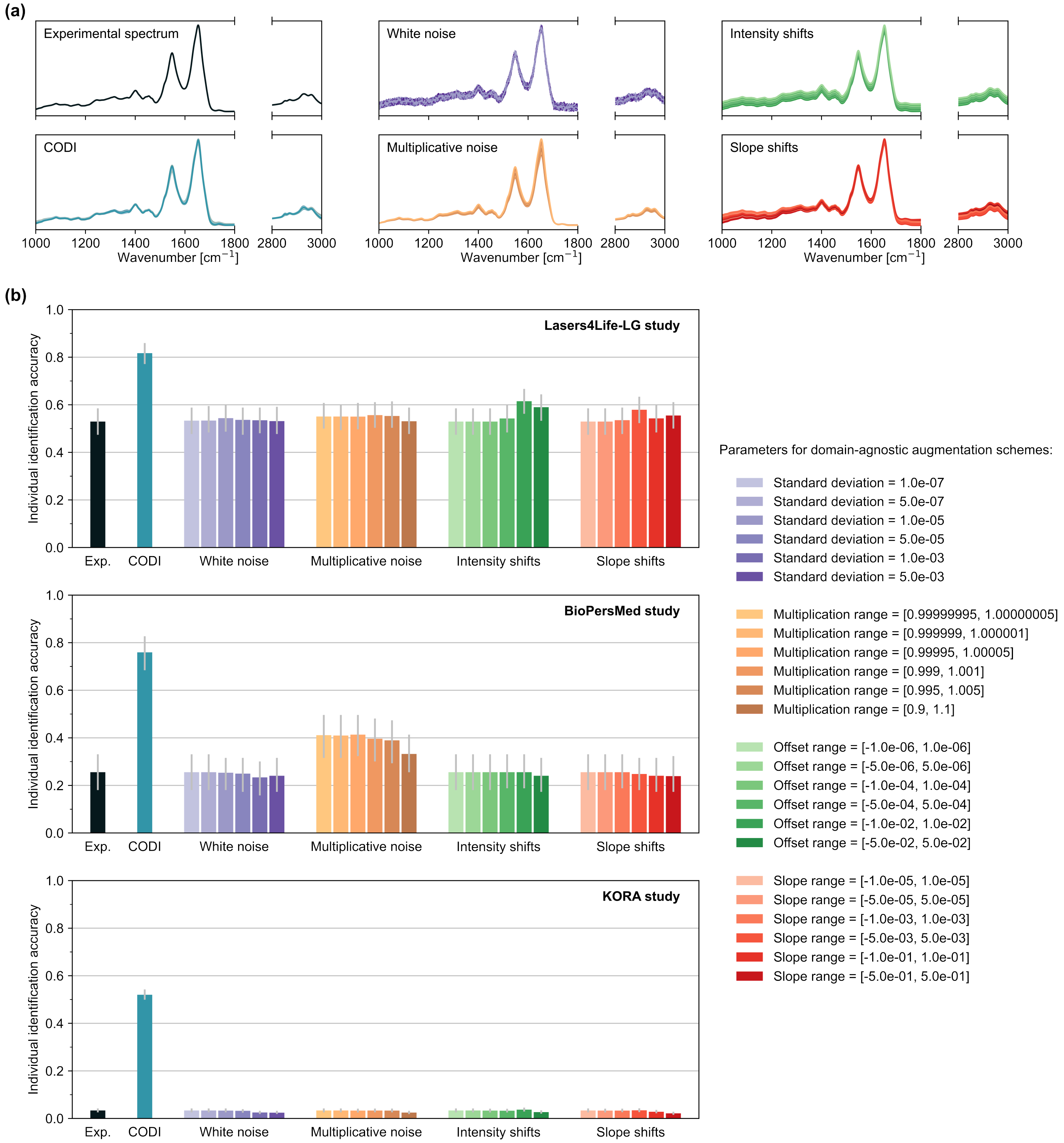


**Figure S2.** Comparison of CODI to domain-agnostic augmentation schemes. (**a**) Given an input seed experimental spectrum (upper left panel), several methods were employed to generate spectra with added variance. These methods include CODI (bottom left panel), as well as other domain-agnostic augmentation schemes. White noise (upper middle panel) was introduced by repeatedly generating a random Gaussian vector and adding it to the input seed. Multiplicative noise (lower middle panel) was introduced by repeatedly scaling the input seed with random scaling factors. Intensity shifts (upper right panel) were introduced to the input seed by vertically shifting the intensity with random factors. Slope shifts (lower right panel) were introduced by repeatedly manipulating the linear slope of the input seed. (**b**) The individual identification classification task was investigated across three study cohorts. The baseline measurement of each individual in the cohort served as the training seed measurements. Classifiers were trained on the experimental data (black bars), data generated through CODI (blue bars), and data generated through the domain-agnostic augmentation schemes (remaining bars). For each domain-agnostic augmentation scheme, several parameters controlling the data generation were examined, as listed in the legend to the right.

[5] T. Eissa, C. Leonardo, K. V. Kepesidis, F. Fleischmann, B. Linkohr, D. Meyer, V. Zoka, M. Huber, L. Voronina, L. Richter, A. Peters, and M. Zigman, "Plasma infrared fingerprinting with machine learning enables single-measurement multi-phenotype health screening," *Cell Reports Medicine*, Accepted Manuscript.

[6] H.-E. Wichmann, C. Gieger, and T. Illig, "KORA-gen - resource for population genetics, controls and a broad spectrum of disease phenotypes," *Das Gesundheitswesen*, vol. 67, pp. 26–30, July 2005.

[7] C. Sujana, J. Seissler, J. Jordan, W. Rathmann, W. Koenig, M. Roden, U. Mansmann, C. Herder, A. Peters, B. Thorand, and C. Then, "Associations of cardiac stress biomarkers with incident type 2 diabetes and changes in glucose metabolism: KORA F4/FF4 study," *Cardiovascular Diabetology*, vol. 19, Oct. 2020.

[8] W. Rathmann, B. Haastert, A. Icks, H. Löwel, C. Meisinger, R. Holle, and G. Giani, "High prevalence of undiagnosed diabetes mellitus in southern germany: Target populations for efficient screening. the KORA survey 2000," *Diabetologia*, vol. 46, pp. 182–189, Feb. 2003.

[9] M. Huber, K. V. Kepesidis, L. Voronina, M. Božić, M. Trubetskov, N. Harbeck, F. Krausz, and M. Žigman, "Stability of person-specific blood-based infrared molecular fingerprints opens up prospects for health monitoring," *Nature Communications*, vol. 12, Mar. 2021.

[10] B. Sanchez-Lengeling, J. N. Wei, B. K. Lee, R. C. Gerkin, A. Aspuru-Guzik, and A. B. Wiltschko, "Machine learning for scent: Learning generalizable perceptual representations of small molecules," *arXiv*, 2019.

[11] E. J. Bjerrum, M. Glahder, and T. Skov, "Data augmentation of spectral data for convolutional neural network (CNN) based deep chemometrics," *arXiv*, 2017.

[12] A. Lebrun, H. Fortin, N. Fontaine, D. Fillion, O. Barbier, and D. Boudreau, "Pushing the limits of surface-enhanced raman spectroscopy (SERS) with deep learning: Identification of multiple species with closely related molecular structures," *Applied Spectroscopy*, vol. 76, p. 609–619, Mar. 2022.
